# Experimental and Computational Analysis of HIFU Thermal Ablation in Breast Cancer Cells: Monolayers vs. Spheroids

**DOI:** 10.1101/2023.12.04.569950

**Authors:** Heba Badawe, Jean Paul Harouz, Kareem Abu, Petra Raad, Kamel Abou Ghali, Wassim Abou Kheir, Massoud Khrariche

## Abstract

**Objective:** The primary objective of our study was to investigate the efficiency of high intensity focused ultrasound (HIFU) ablation in two distinct cellular configurations, 2D monolayers and 3D spheroids of epithelial breast cancer cell lines. The study also compares empirical findings from experiments with results obtained through numerical simulations using a bioheat computational model. This comparison is intended to provide a comprehensive understanding of the acoustic energy conversion within the biological system during HIFU treatment.

**Methods:** HIFU was applied to 2D and 3D cultured MDA-MB 231 and MCF7 epithelial breast cancer cell lines while systematically varying ultrasound intensity and duty cycle (DC) during sonication sessions of different durations. Temperature elevation was measured and the ablation percentage was calculated based on bright field and fluorescent imaging of the treated regions. Experimental results were validated through simulations of the ablation setup.

**Results:** Upon HIFU, spheroids exhibited a lower temperature increase (approximately 20 °C) when subjected to comparable acoustic intensities and duty cycles. The level of tumor ablation was highly influenced by DC, with higher DCs leading to greater ablation percentages. However, sonication duration had a minimal impact on the degree of ablation. Numerical simulations corroborated these observations, demonstrating uniform heat distribution within the cultured cells. At higher DCs and intensities, complete ablation of spheroids was achieved, whereas at lower levels, only the outermost layers exhibited ablation.

**Conclusion:** Our study reveals a significant disparity in the response of 2D monolayers and 3D spheroids to HIFU treatment. Specifically, tumor spheroids require lower temperature elevations for effective ablation, and their ablation percentage significantly increases with elevated DC.

## 1. Introduction

Cancer poses a significant global public health challenge and currently ranks as the second leading cause of death worldwide, following cardiovascular diseases. Notably, among women, breast cancer emerges as the most extensively studied malignancy and a foremost contributor to cancer-related fatalities (1, 2). Traditional treatments for breast cancer, on the other hand, encompass a range of modalities contingent on factors such as cancer type, treatment tolerability, and disease stage. These conventional screening and treatment methodologies encompass invasive surgery, immunotherapy, chemotherapy, and radiotherapy, each carrying notable risks to patient health and well-being (3, 4). These interventions often entail surgical procedures designed to excise tumors, or the administration of chemotherapeutic agents via intravenous infusion, targeting rapidly proliferating cancer cells (5). However, the invasiveness of these approaches, though effective to some degree, gives rise to a host of adverse effects. These can include imprecise delineation of the tumor leading to damage to adjacent tissues (6), complications stemming from drug administration, hemorrhaging, infections, substantial illness (7) and protracted post-treatment recovery requiring extended hospitalization (8, 9).

High-Intensity Focused Ultrasound (HIFU) has emerged as an alternative, non-invasive, and non-ionizing approach for precisely ablating tumor cells. HIFU can generate coagulative necrosis within a specific focal area within the body while sparing adjacent structures along the path of the acoustic beams (10, 11). In general, HIFU induces two major biological effects on cell ablation: thermal and non-thermal (2, 12, 13). Thermal ablation results from the conversion of acoustic energy to thermal energy at the target site through an increase in the energy density, thus elevating the temperature of the cells to a range of 60–85 °C within seconds (14). These elevated temperatures within the focal zone initiate protein coagulation and cell membrane fusion, ultimately leading to the necrosis of tumor cells. Beyond this focal region, heat diffusion generates a temperature gradient, where cells do not experience an instantaneous fatal thermal dose but are subjected to temperatures exceeding 40 °C. In contrast, the mechanical or non-thermal ablation involves the transfer of acoustic power by HIFU waves, inducing cavitation at the target site and resulting in the mechanical disruption of cell membranes. High-pressure acoustic waves induce changes in the gaseous composition within tissues, instigating oscillation and subsequent rupture of bubbles, causing subcellular-scale mechanical damage to tissues (15, 16).

HIFU has been successfully employed to treat patients having cancerous tumors in the liver, breast, kidney, and pancreas (17, 18). Yet, this technology still requires adequate research and study. Its efficacy in tumor ablation relies on a multitude of factors, contingent upon the ultrasound parameters employed and the specific target areas, all of which significantly influence its success rate. Consequently, there is a pressing imperative for comprehensive exploration and refinement of HIFU technology to optimize therapeutic outcomes. Enhancing the efficiency of HIFU tumor treatment necessitates thorough in vitro assessments, involving the cultivation of tumor cells for preclinical cancer research. Traditional cell culturing methods involve the growth of cells in monolayers, which replicate along a single plane. The advantages of this 2D cell culture model include its simplicity, cost-effectiveness, and ease of maintenance (13), making it suitable for basic cancer studies. Nevertheless, this two-dimensional cell model fails to replicate the intricate cell-cell and cell-extracellular environment interactions crucial for a comprehensive understanding of cells’ natural behavior in vivo. These interactions encompass processes such as differentiation, proliferation, gene and protein expression, and cellular responses to external stimuli (19).

To overcome the limitations inherent in the two-dimensional model, investigators turn to spheroid models. Spheroids offer a three-dimensional configuration that permits cells to interact and proliferate in all directions, fostering the development of an extracellular architecture with a layered structure and a proliferative profile (20). This 3D environment accounts for the presence of oxygen and nutrients, allowing cells to grow with a gene expression profile closely resembling that of tumor cells. Consequently, the use of spheroid 3D cell configurations offers a more accurate evaluation of biological responses compared to the conventional 2D monolayer cell formation (21).

In this study, we investigate the use of HIFU for in vitro thermal ablation of epithelial breast cancer cell lines. These cell lines were cultured in either 2D monolayer or 3D spheroidal configurations. The study includes both experimental and computational analyses of how these two cell cultures interact with acoustic waves. By utilizing these two distinct geometrical models, we emphasize that 3D spheroids provide a more representative representation of tumor cells undergoing ultrasonic ablation. We studies ultrasound parameters, specifically acoustic intensity, duty cycle (DC), and sonication duration (SD), and assessed their impact on temperature elevation and the extent of tumor cell ablation. Furthermore, we compared our empirical findings with the results from numerical simulations using our adopted bioheat computational model. This model predicts the conversion of acoustic energy within the biological system.

## 2. Materials and Methods

### 2.1 HIFU Mapping and Characterization

Characterizing the ultrasound transducer required determining its power output per unit volume and focal point. The utilized transducer of a 2 MHz center frequency (SU101-019; Sonic Concepts) generated HIFU waves after receiving a sinusoidal electrical signal from a function generator (SIGILENT-SDS 1025), pre-amplified by a 50-ohm RF power amplifier (100L Broadband Power Amplifier, Electronics & Innovation Ltd). Generally, the transducer converted the electrical signals into propagating mechanical ultrasonic waves that were captured by an immersible needle hydrophone (Onda HNR-0500). The latter converted them back into voltage recordings to be stored in a data acquisition system, followed by processing and quantizing into acoustic intensity (22). The hydrophone was originally mounted on a 3D-motorized axes system, allowing its systematic movement into different positions relative to the transducer to scan a definite geometric volume (Figure 1A). Using this transmission method, the focal point of the ultrasound transducer, where the maximum intensity of 5 𝑊/𝑐𝑚^2^was recorded, was localized at 51.2 ± 0.1 mm from the face of the transducer in the free-field tank filled with degassed, distilled water (Figure 1B).

**Figure 1.**
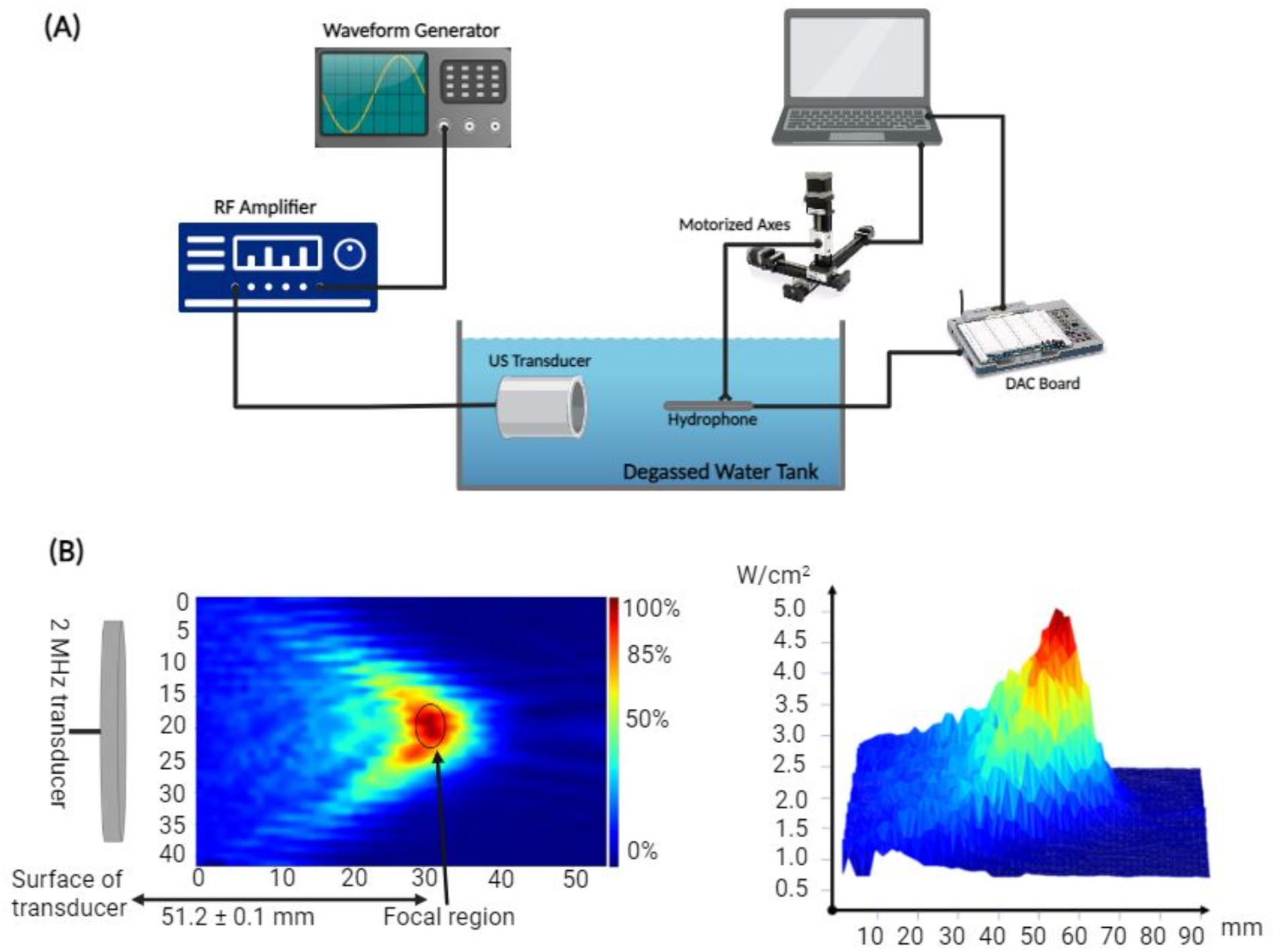
Acoustic profile setup. (A) Schematic diagram of the experimental setup used to determine the acoustic intensity of HIFU emitted by the transducer and the temperature rise at target. (B) Heat map displaying the intensity distribution at the focal point of the transducer present 51.2 mm away from the center of curvature of its surface.

### 2.2 Temperature Acquisition

For temperature measurements in the focal region of the ultrasound transducer, used in the 2D and 3D cell ablation experiments, a K-type thermocouple (Omega) was taped in place over a gel-filled glass slide placed at the focal point within a petri dish (Figure 2A). The glass slide could withstand very high temperatures. For the delivery of ultrasound waves, a coupling cone, sealed with an acoustically transparent membrane, was placed over the inverted ultrasound probe so that the focal area was within 1 mm from the tip of the probe. To cool the transducer off and remove any potential gas bubbles that may form within the cone, deionized water was continuously pumped in and out of the cone using a pump-perfusion kit (PPS2, Multichannel systems) at a maximum flow rate of 30 mL/min. The cone was then coupled to the petri dish containing the glass slide. An IR-FLIR E40 Thermal Imaging Camera was also vertically mounted over the petri dish (Figure 2B) to collect quantitative data for the examination of any temperature variation within and outside the focal region of the focused ultrasonic transducer.

**Figure 2.**
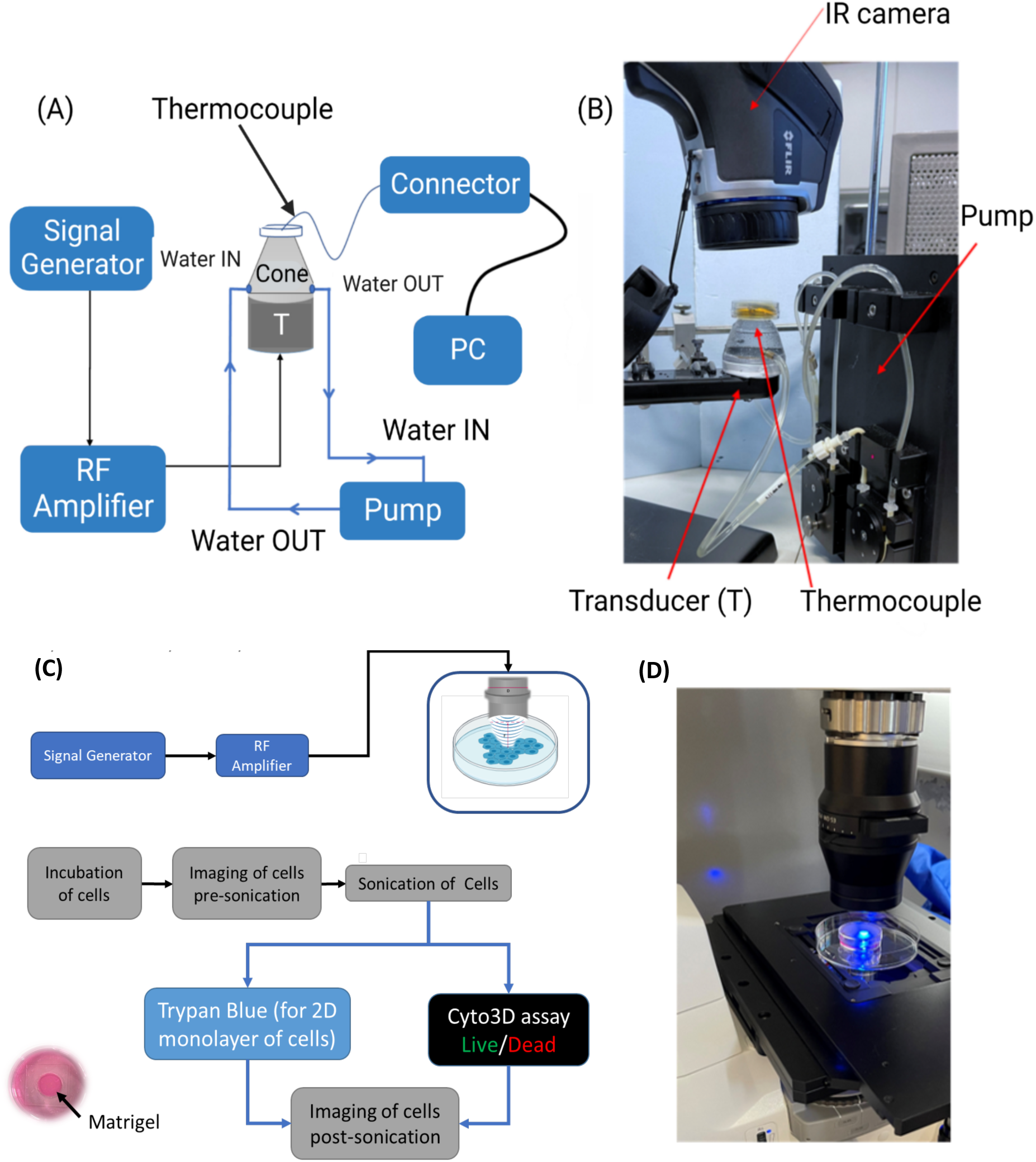
Temperature measurement at the focus and ultrasound ablation setup. (A) Schematic diagram of the setup to measure the temperature. The thermocouple was inserted into the gel-filled glass to record the maximum temperature at the end of sonication time. (B) Photo of the experimental setup, the temperature at and around the focus was captured by the IR camera. (C) Schematic diagram of the experimental setup where cultured tumor cells are treated with HIFU at distinct sets of ultrasound parameters and then stained and visualized under the microscope. (D) Photo of the petri dish in which cultured cells are being stained after HIFU treatment to determine the ablation percentage.

### 2.3 2D Tumor Cell Culturing

MDA-MB 231 and MCF7 epithelial breast cancer cell lines were considered for this study. The cells were cultured in a glucose-rich DMEM medium containing 10% fetal bovine serum (FBS) (Sigma, F-9665), and 1 % penicillin/streptomycin (Lonza, Basel, Switzerland, DE16–602E) with 5 % carbon dioxide in a humidified incubator, at a temperature of 37 °C(23). Such medium was rich in nutrients, growth factors, and antibiotics, where the FBS provided a source of proteins, and the penicillin/streptomycin helped in the prevention of any bacterial contamination. The medium was replenished after 36 hours to remove any waste products. Around 3,000 cells were plated on sterile coverslips and incubated for 24-36 hours before ultrasound ablation. The cells were ready for treatment after reaching a confluence of more than 95 %. Throughout the process, from seeding to treatment, the cells were checked under an inverted bright field microscope to ensure healthy proliferation and absence of contamination(24).

### 2.4 3D Tumor Cell Culturing

The characteristics of sphere-like MCF-7 and MDA-MB 231 breast cancer cells were studied using a sphere formation assay. In this assay, 3D tumor spheroids that mimic the environment of the extracellular matrix (ECM) were formed. The single cells were mixed with a 1:1 combination of cold Matrigel (a growth factor-reduced substance) and serum-free StableCellTM RPMI-1640 at a density of 5000 cells/dish. The mixture was poured as droplets in microwells present in the center of small petri dishes and allowed to solidify at 37 °C in a humidified incubator with 5 % CO2 for 1 hour. Afterward, 1.5 mL of StableCell TM RPMI-1640 cell growth medium containing 5 % FBS was added to each petri dish. The medium was replaced every 2-3 days until the spheroids matured and were ready for further experimentation, specifically ultrasound sonication. The cells were incubated for a total of 9 days while maintaining a consistent incubation period across all experiments to ensure consistency.

### 2.5 Ultrasound Cellular Ablation

Before any treatment, the cells were examined in each petri dish to ensure even growth, and pictures were taken using a bright field microscope. To measure the ablation area, a glass slide with nine small squares, each measuring 100x100 µm^2^, was placed under the cells. These squares were positioned so that one covered the middle of the focal region while the other eight covered spots scattered around and outside the treatment zone. Ultrasound sonication of the cultured cells was carried out using an ultrasound transducer coupled with a cone and aligned with the middle of the glass slides containing the cells (Figure 2C). Bright field and fluorescent images of cellular ablation were taken at different magnifications post ultrasound sonication (Figure 2D).

### 2.6 Cell Viability Assessment

Post ultrasound sonication, cellular viability was determined using two staining techniques: the Trypan Blue Exclusion and the Fluorescent Cyto3D Live/Dead Assay. Trypan Blue, a straightforward and user-friendly stain, was utilized. Upon application, it rendered living cells unstained, while nonviable cells appeared distinctly blue when observed under a microscope. The Fluorescent Cyto3D Live/Dead Assay is a non-toxic nucleic dye which marks damaged cancer cells in red using Acridine Orange (AO), and alive cancer cells in green with Propidium Iodide (25, 26). This dye was applied before and after treating the tumor cells with ultrasound sonication. For both the 2D and 3D cell cultures, the dye had to be mixed with pre-prepared media containing 2 µL of Cyto3D for every 100 µL of media. This mixture was prepared in a sterile, light-sensitive environment conditions to prevent any signal loss. After completely removing the media from the petri dish, 600 µL of the prepared solution was added directly to the Matrigel and incubated for at least an hour to allow the chemicals to permeate through the gel and into the cell colonies and spheres.

For cell visualization, an inverted microscope was employed to capture bright field and/or fluorescent images. Cells were also examined for viability pre-treatment to guarantee that cell death was solely due to the ultrasound treatment. Cell quantification was conducted on 3D Matrigel cultures using the following procedure: Initially, all the culture medium was completely withdrawn and transferred to a conical tube, which already contained previously collected media from the petri dish. Next, cold trypsin was introduced to the petri dish and allowed to act for approximately 5 minutes. Following that, the cells were gathered and displaced from the dish to the same conical tube. After thoroughly mixing the contents, 50 µL were drawn from the conical tube and combined with 50 µL of Trypan Blue. Finally, 10 µL of the resultant mixture were loaded onto the hemocytometer for the purpose of cell counting.

### 2.7 Statistical Analysis

Ablation data were obtained across four spatial-peak pulse-average intensities covering 146.7 W/cm^2^ to 500 W/cm^2^ for each DC spanning from 15 % to 55 %. The ablation data were analyzed with non-parametric tests using IBM SPSS Statistics (29^th^ version) given the small sample size per DC and where the data were not normally distributed. Multiple pairwise comparisons were conducted utilizing Kruskal Wallis test and Dune-Bonferroni corrected to determine significant differences in cell ablation among different DCs for a specified SD. Additionally, within a particular DC, two independent ablation samples from different SDs were compared using the Mann-Whitney U-test. Combining all duty cycles, the Mann-Whitney U-test was used to assess differences in ablation levels at the focal area caused by SDs of 5 minutes and 10 minutes. Statistical significance was determined at p < 0.05.

### 2.8 Computational Modeling

We developed mathematical models to assess the thermal response of both monolayer and spheroid cell cultures to pulsed HIFU acoustic sonication for results validation and outcome predictions. The models were validated with experimental data and used to conduct a parametric study for the different operating conditions such as DC and maximum pressure to determine the effect of HIFU on cellular ablation. The wave propagation model provided the acoustic field distribution and the corresponding thermal energy deposition in the cells. The thermal energy was used in the thermal model to determine the temperature distribution in the cell cultures and their corresponding ablation rate.

#### 2.8.1 Computational Domain

The developed model was used to determine the temperature distribution inside the cell cultures of both configurations, monolayer and spheroids. For this reason, the adopted computational domain, presented in Figure 3A, consisted of a petri dish and a transducer surface. The petri dish in each configuration was formed of different layers, starting from the transducer surface as follows: cooling water, cell growth medium, cells, glass, and the petri dish bottom surface. To simplify the computational domain for the spheroids cell culture, it was assumed that they occupy a homogenous layer, neglecting the voids created between the different hydrogel spheres where the cells were seeded. This could be adopted due to the relatively small (∼μm) dimensions of those spheres. Moreover, the cooling water was separated from the cell culture by a thin polymeric membrane that was disregarded in the computations since it is basically acoustically transparent (27).

**Figure 3.**
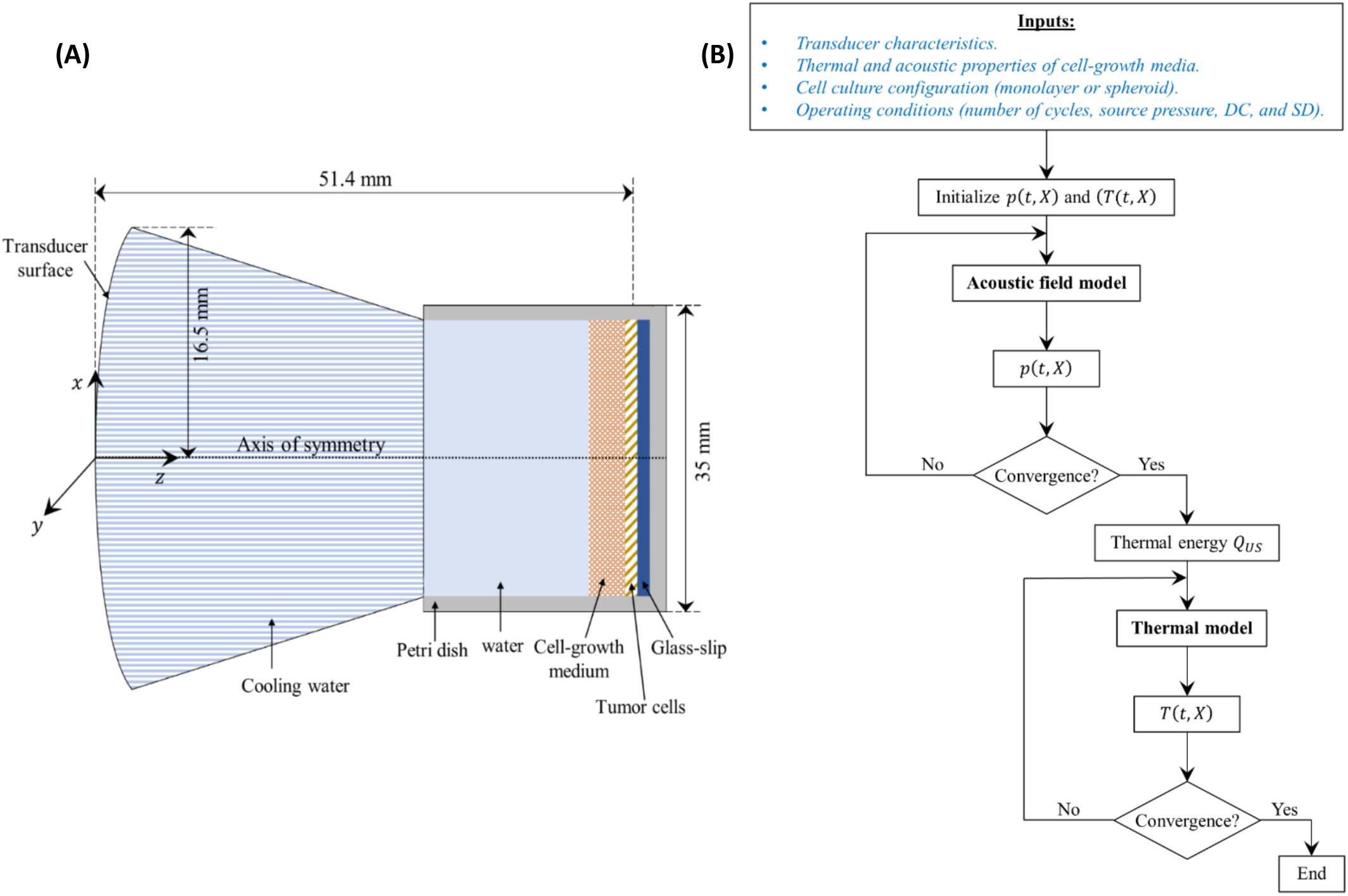
Computational model. (A) Schematic diagram of the computational domain adopted for the numerical simulations. (B) Adopted solution flowchart for the developed mathematical models.

#### 2.8.2 Acoustic Field Model

Ultrasound waves emitted from the transducer traveled through the cooling water before passing through the culture medium where the cells were grown (28). At the interface of each layer, part of the acoustic energy was absorbed while the remaining energy was reflected. The fraction of reflected energy depended on the difference in the various layers of acoustic impedance (Z (kg/𝑚^2^·s)), which is the product of the layer’s density (ρ (kg/𝑚^2^)) and the acoustic wave velocity (𝑐_o_ (m/s)) (29). Within a single layer, the pressure fluctuations caused shearing of the cells, resulting in increased mechanical friction that was converted to thermal energy (30, 31).

To model the wave propagation inside the cell cultures, the acoustic pressure field 𝑝(𝑡, 𝑋)(Pa) was determined (32). The Westervelt equation was adopted as it provided a more general acoustic field model to the commonly used Khokhlov–Zabozotskaya– Kuznetsov (KZK) equation (33). The Westervelt equation is derived from the fluid motion equations, making it more accurate than the KZK equation which suffers from a validity region (34). Such a model has been commonly adopted in medical ultrasound as it combines the influence of absorption, non-linearity, and diffraction mechanisms that are present in biological media (35) and given by:

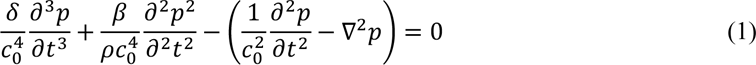

The first term of equation (1) describes the loss caused by the viscosity and heat conduction in the fluid. The second term describes the nonlinear distortion of the propagation wave, and the last two terms represent the linear lossless wave propagation. The different terms of the equation represent the following: ∇^2^ (m^-2^) is the Laplace operator, 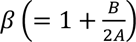 is the coefficient of nonlinearity, which is a function of the nonlinearity parameter 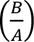 of the medium. Acoustic diffusivity is represented by 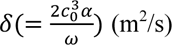 which depends on the angular frequency (𝜔 (rad/s)) of the source and the absorption/attenuation coefficient (𝛼 (Np/m)). The latter is determined by the frequency-dependent power law given by (36):

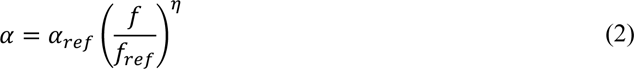

where 𝛼*_ref_* (Np/m) is the attenuation coefficient at a reference frequency 𝑓*_ref_* of 1 MHz and η is the attenuation exponent of the power law and it depends on the type of the medium (37).

Once the acoustic pressure 𝑝(𝑡, 𝑋) was determined at each point 𝑋(𝑥, 𝑦, 𝑧) of the cell culture domain, the resulting heat (thermal energy) deposition from the ultrasound wave (𝑄*_US_* (W/m^3^)) was computed from the acoustic intensity (𝐼 (W/m^2^)) and the medium impedance (Z) using equation 3:

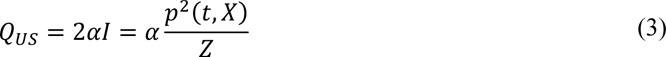

The model entailed an initial condition of acoustic pressure, which was set to zero at the beginning of sonication. Additionally, a set of boundary conditions was required to solve for the acoustic field. The geometry of the petri dish enabled an axisymmetric solution around the z-axis, reducing the computational domain. Hence, a symmetry boundary condition was adopted at planes x=y=0. The transducer emitted a pulsed HIFU beam at z=0 (38), which was expressed by:

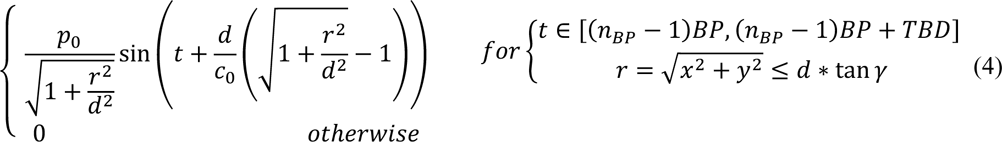

where 𝑝_o_(Pa) is the pressure at the transducer surface, 𝑛*_BP_* is the number of burst cycles with a period of TBD (s) each during the total sonication duration. r (m) is the radial distance from the center of the transducer with a focal distance d (m), within the plane 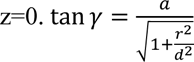, with γ (rad), is the aperture angle which is a function of the transducer radius 𝑎 (m) and the focal distance d (m).

Finally, since the computational domain was finite, the perfectly matched layer boundary condition was adopted at the bottom of the petri dish (end surface) to control the acoustic reflections from the end of the domain using artificial absorption (39).

#### 2.8.3 Thermal Model

Once the acoustic pressure was established in the petri dish, the temperature distribution could then be determined from the thermal model. The energy balance presented in the bioheat model of Pennes (40) was commonly used in these applications (32). The model was based on the Fourier law, and considered the effect of blood flow in biological media as well as the metabolic heat generation (𝑄_5_ (W/m^3^)). The temperature distribution ((𝑇(𝑡, 𝑋) (K)) was thus computed from equation (5):

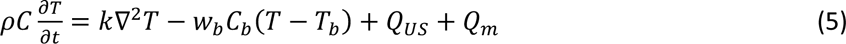

where C (J/kg·K) and k (W/m·K) are the respective specific heat capacity and thermal conductivity of the medium. 𝐶_6_ (J/kg·K), 𝑤_6_ (kg/s·m^3^) and 𝑇_6_ (K) are the specific heat capacity, perfusion rate and temperature of the blood respectively.

For the cell culture in the petri dish, the effect of blood perfusion was omitted due to the absence of blood vessels in the culture-growing medium. Moreover, the metabolic heat generation could be neglected due to its small contribution to the overall energy balance as compared to the thermal energy from the acoustic wave (32). To solve the bioheat equation, the initial temperature was set to 37 °C. In addition, symmetry boundary conditions were considered for planes x=y=0, while Dirichlet boundary conditions of 25 °C were set for both end surfaces of the domain(41).

#### 2.8.4 Numerical Solution

The mathematical models of the wave propagation and temperature distribution were solved using the finite volume method with implicit Backward Euler temporal scheme and central difference scheme for second order spatial differentials. The nonlinear and absorption terms of equation (1) were solved as presented by Doinikov et al. (34) for a transient 3D Cartesian grid. To ensure accuracy of the results with minimal computational time, the time step was set to Δt=0.01/𝑓_0_ with a grid of Δx=Δy=0.2λ and Δz=0.1λ (42), where λ (=𝑐_0_/𝑓_0_) (m) is the wavelength of the driving frequency of the transducer. Once a uniform pressure field was obtained at steady state (no change in the acoustic pressure with time), the acoustic intensity and the corresponding thermal energy generated in the cell cultures were determined and used in the thermal model with a time step of Δt=0.01 s. Different time steps were adopted for the thermal model compared to the acoustic field model to reduce the computational time (32). The convergence of the different parameters (𝑝(𝑡, 𝑋), 𝑇(𝑡, 𝑋)) was reached when their calculated residuals between two consecutive iterations were less than 10^-8^. The different thermal and acoustic properties of the monolayer and spheroid cell cultures, as well as the different layers through which the acoustic wave was propagating, are presented in Table 1. Note that the missing acoustic parameters for either the cells or their growth medium were adopted similar to those of water as the case for most biological tissues (29, 43).

**Table 1.**
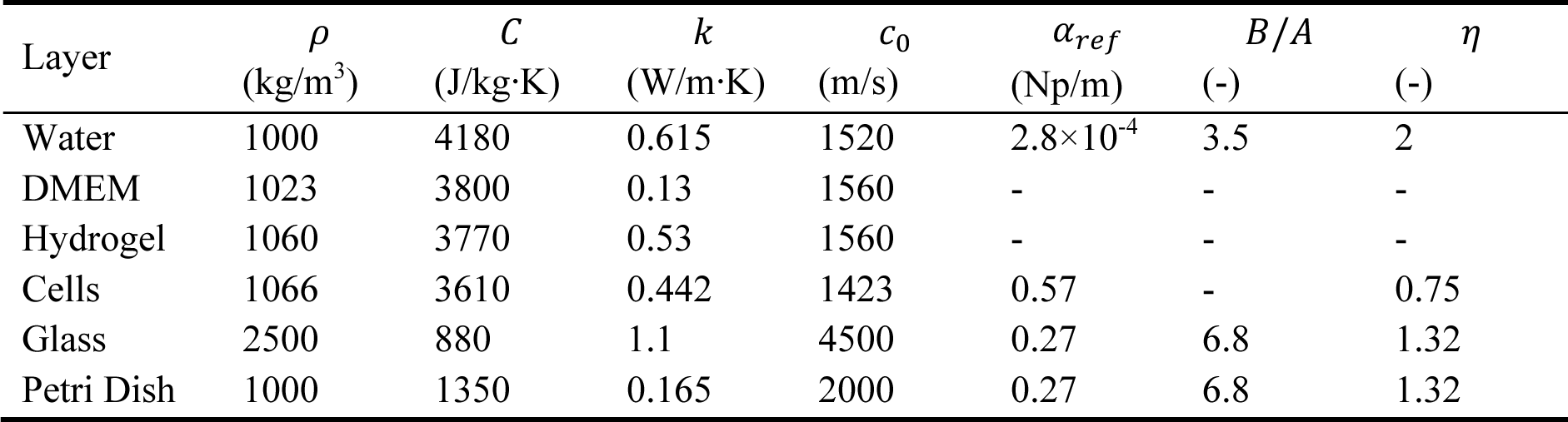
Thermal and acoustic properties of the different layers of the cell cultures.

| Layer | $\rho$<br>(kg/m<sup>3</sup>) | $C$<br>(J/kg·K) | $k$<br>(W/m·K) | $c_0$<br>(m/s) | $\alpha_{ref}$<br>(Np/m) | $B/A$<br>(-) | $\eta$<br>(-) |
| --- | --- | --- | --- | --- | --- | --- | --- |
| Water | 1000 | 4180 | 0.615 | 1520 | $2.8 \times 10^{-4}$ | 3.5 | 2 |
| DMEM | 1023 | 3800 | 0.13 | 1560 | - | - | - |
| Hydrogel | 1060 | 3770 | 0.53 | 1560 | - | - | - |
| Cells | 1066 | 3610 | 0.442 | 1423 | 0.57 | - | 0.75 |
| Glass | 2500 | 880 | 1.1 | 4500 | 0.27 | 6.8 | 1.32 |
| Petri Dish | 1000 | 1350 | 0.165 | 2000 | 0.27 | 6.8 | 1.32 |

The solution of the developed mathematical model followed the flowchart presented in Figure 3B. The model inputs included the transducer characteristics (a, d, 𝑓_0_, 𝑝_0_), the cell cultures thermal and acoustic properties and configuration (monolayer, spheroid) and the operating conditions (TBD, DC, SD). It started by initializing the pressure and temperature fields and used the inputs to determine the converged steady state acoustic pressure distribution from the acoustic field model. The latter was incorporated to determine the thermal energy deposition (𝑄*_US_*) that was inputted in the thermal model to conclude the converged temperature distribution.

## 3. Results

### 3.1 Focal Area and Temperature Measurements

The focal area at the horizontal mid-plane varied as a function of increasing input voltage. When using a 5 MHz focused ultrasound transducer, the focal area increased from 3.5 mm^2^ to 5.2 mm^2^ as the input voltage increased from 110 mV to 160 mV (Figure 4A). Temperature measurements were acquired in real-time using a thread-head K-type thermocouple set at the focal area of the ultrasound transducer while sonicating at different voltages and duty cycles. While varying the input voltage, the temperature at the focal area increased as well with an increase in the duty cycle of ultrasound waves (Figure 4B). The increase was monotonic, recording a minimum temperature of 43 °C at the lowest input voltage of 300 mV and a DC of 10 %, reaching the maximum temperature of 81.9 °C, when sonicating at an input voltage of 350 mV at the highest DC of 45 %. The IR camera images emphasized a uniform distribution of thermal energy throughout the focal region of a recorded 134.5 mm^2^ area when sonicating with ultrasound waves of 10 % DC and an input voltage of 350 mV.

**Figure 4.**
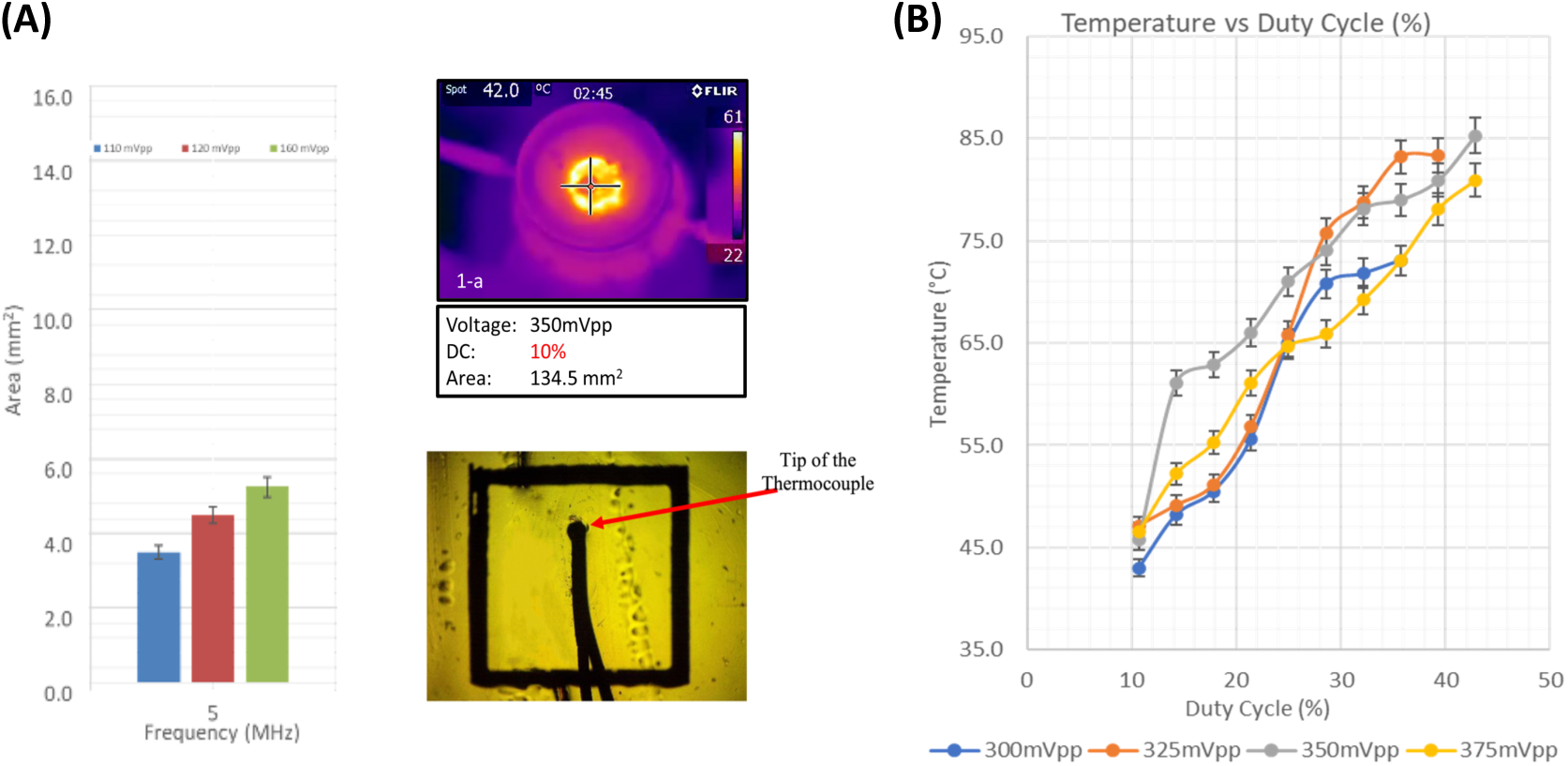
Focal area and temperature measurements. (A) Focal area as a function of amplitude of input signal. Heat distribution at the focus as captured by the IR camera and a close up photo of the thermocouple in position. (B) Temperature measured by the thermocouple as a function of duty cycle at distinct input voltage amplitudes.

### 3.2 Cellular Ablation Using HIFU

#### 3.2.1 On 2D Cultures

Ultrasound sonication was targeted toward a 2D layer of cells seeded on a glass coverslip, taking the configuration of a monolayer of cultured cancer cells. HIFU of frequency 2 MHz at a 35 % duty cycle, and 280 𝑊/𝑐𝑚^2^ intensity caused a 95 % ablation of the 2D cultured cells inside the focal region, as opposed to a negligible 3 % ablation outside the focal region. Bright-field images of the cultured cells before and after ultrasound sonication for 10 min emphasized the difference in the ablation percentage (Figure 5A), with a maximum temperature of 60.9°C reached inside the area of focus, approximately 1.5 times higher than that achieved outside the transducer’s focus (Figure 5B).

**Figure 5.**
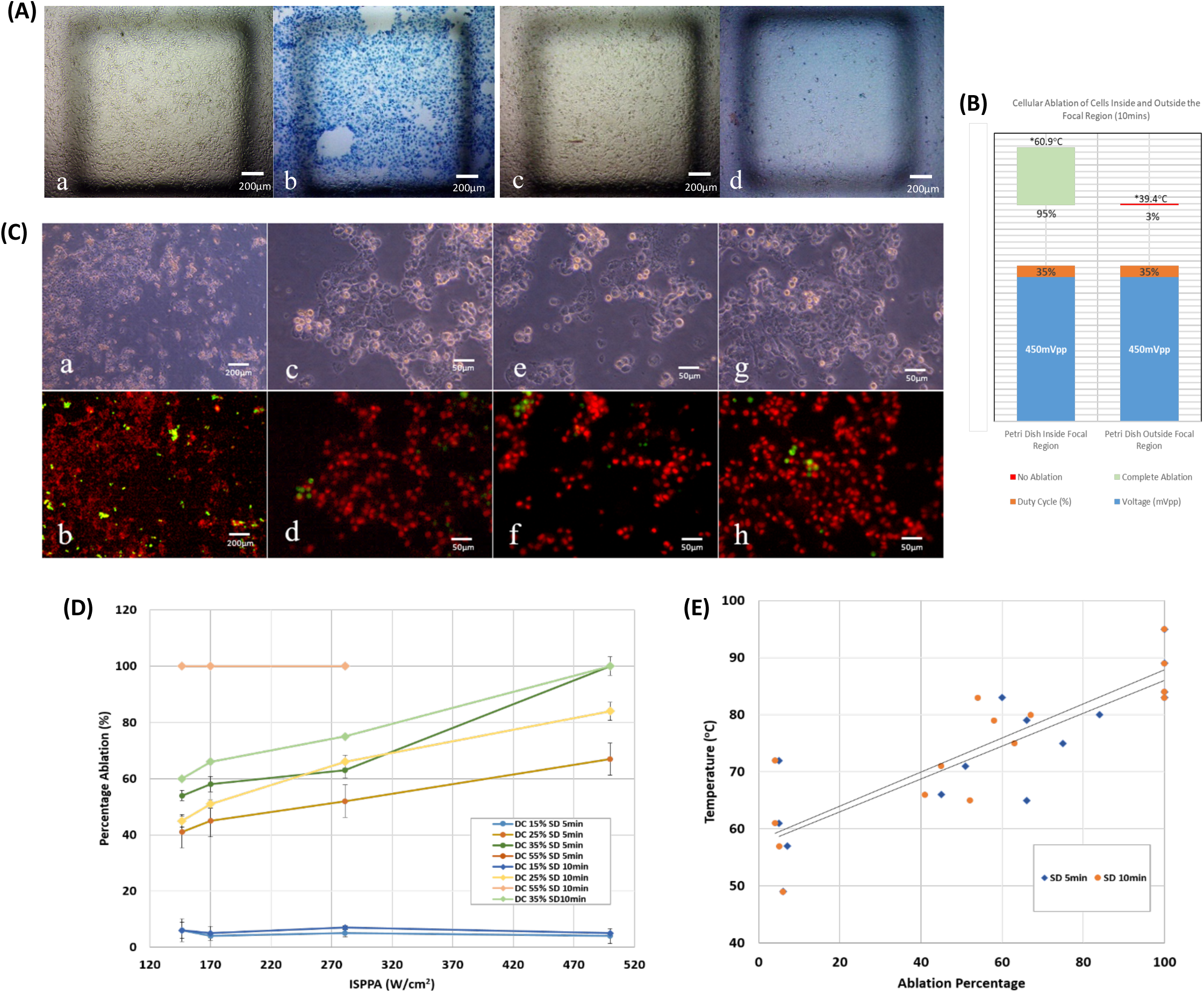
Monolayer ultrasonic ablation. (A) Bright field images, (a & b) cells located inside the focal region, pre-sonication and post-sonication respectively, with dead cells stained in blue. (c & d) cells located outside the focal region, pre-sonication and post-sonication respectively. (B) Ablation percentage following 10 minutes of sonication. The ablation area is greater than 95 % inside the focal region, whereas it is in the vicinity of zero outside the focal region. (C) Bright field images of cells post sonication at (a) 5x magnification and at 20x magnification (c, e, g). Fluorescent images of cells at (b) 5x magnification and at 20x magnification (d, f, h). Results show 99 % ablation where alive cells are stained in green, while dead cells are stained in red. (D) Percentage ablation of monolayers as a function of acoustic intensity at distinct DC and SD. (DC =15, 25, 35 or 55 %, SD=5 or 10 minutes. (E) Final temperature versus ablation percentage of monolayers following ultrasound sonication for 5 or 10 minutes.

Sonication duration and duty cycle of ultrasound waves affected the ablation percentage as a function of the applied spatial peak pulse average ultrasonic intensity (Figure 5D). Regardless of the sonication duration, a 15 % low duty cycle and a 55 % high duty cycle caused overlapping in the ablation percentage, which constantly varied as 𝐼*_SPPA_* increased. However, all cells got ablated at a 55 % DC, yet approximately less than 5 % got ablated at a 15 % DC. In general, incrementing the sonication duration did not necessarily cause a significant change in ablation, contrary to increasing the duty cycle and 𝐼*_SSPA_* At a 35 % DC and at the highest 𝐼*_SPPA_* of 500 𝑊/𝑐𝑚^2^, full ablation of cultured tumor cells occurred (Figure 5C). With the exception of the highest DC of 55 %, a low 𝐼*_SPPA_* (≤ 170 W/cm^2^) could only achieve an ablation percentage less than 70 % with all applied DC and SD.

Majorly, thermal ablation of cancerous cells cultured in 2D started at temperatures around 60 °C to reach full ablation at temperatures above 80 °C, regardless of the sonication time applied (Figure 5E). The ultrasound treatment time had a negligible effect on the temperature and ablation levels reached (p=0.557). The percentage of ablated 2D cultured cells increased monotonically with an increase in the temperature.

#### 3.2.2 On 3D Cultures

Cancerous cells 3D cultured in Matrigel received ultrasound sonication at a frequency of 2 MHz, with varying ultrasound intensity, duty cycle, and sonication duration. Bright field images and Fluorescent staining of cells after sonication for 10 min with ultrasound waves of 35 % DC, at an intensity of 280 𝑊/𝑐𝑚^2^, showed a significant 90 % ablation percentage of tumor cells inside the focal region with an elevation in temperature to 61.4 °C, in comparison to a minimal 4% ablation percentage of cells recorded outside the field of focus with an approximate 1.57 times lower temperature reached (Figure 6B). Bright-field images and fluorescent images at 20x magnification showed clusters of dead cells post-sonication reaching up to 99 % ablation at a high DC of 55 % (Figure 6A).

**Figure 6.**
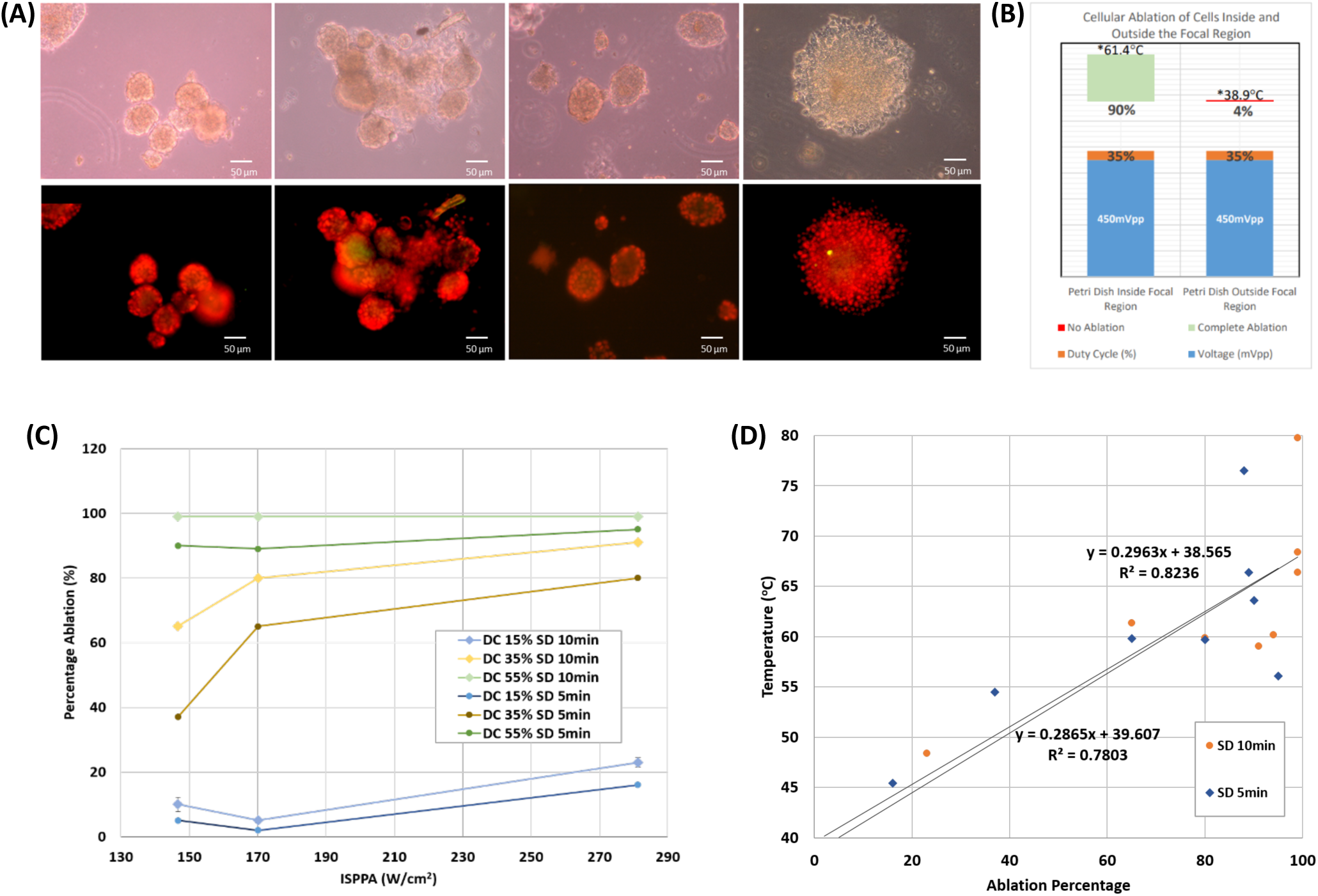
Spheroid ultrasonic ablation. (A) Bright field images of cells post sonication at 20x magnification (upper row). Fluorescent images of cells post sonication at 20x magnification (lower row). (B) Ablation percentage following 10 minutes of sonication. The ablation area is greater than 90 % inside the focal region, whereas it is in the vicinity of zero outside the focal region. (C) Percentage ablation of spheroids as a function of acoustic intensity at distinct DC and SD, (DC =15, 25, 35 or 55 %, SD=5 or 10 minutes. (D) Final temperature versus ablation percentage of spheroids following ultrasound sonication for 5 or 10 minutes.

The time of ultrasound treatment affected the ablation percentage as a function of DC and 𝐼*_SPPA_* as well. At all parameters used, an SD of 10 min caused higher ablation percentages than an SD of 5 min. Full ablation of cancerous cells 3D cultured in Matrigel occurred at the highest DC used of 55 %, after ultrasound sonication for 10 min, regardless of the applied ultrasound intensity (Figure 6C). A low 15 % DC caused very minimal ablation (less than 20 %) even after increasing SD and 𝐼*_SPPA_* In general, upon incrementing DC, SD, and 𝐼*_SPPA_* the percentage of ablated tumor cells increased, with a minimal dependence on SD at low and high DCs, in contrast with DCs in the middle range (35 %) where spheroids reached higher ablation percentages as a function of both SD and 𝐼*_SPPA_*

At the end of each ultrasound sonication, 65 % of the trials ablated cancerous spheroids at a temperature ≥ 60 °C (figure 6D), reaching even higher ablation percentages and final temperatures with an increase in sonication time. Minimal ablation of spheroids was recorded at temperatures starting around 45 °C, following a monotonically increasing trend as the temperature increased.

### 3.3 Spheroid vs Monolayer Thermal Culture Ablation

With the monolayer and spheroidal configurations, increasing DC significantly raised the percentage of ablated cells. A DC of 15 % was the least efficient whereas a DC of 55% caused the ablation of more than 90 % of the cells, even at the lowest intensity. This was the outcome for both the monolayers (55 % DC versus 15 % DC: p = 0.008 for both SD = 5 min and 10 min) and spheroids (55 % DC versus 15 % DC: p = 0.022 and 0.028 for SD = 5 and 10 minutes respectively). However, as long as SD was considered, and in both configurations, similar ablation percentages were achieved with no significant difference at a given DC and 𝐼*_SPPA_* The Mann-Whitney U-test yielded insignificant results when comparing ablation percentages at an SD of 5 min to an SD of 10 min, at a DC of 15 % (monolayers: p = 0.2; spheroid: p = 0.4), and at a DC of 55 % (monolayers: p = 1.0; spheroids: p = 0.2).

Ultrasound sonication of tumor cells having the 2D monolayer configuration, depicted their ablation at temperatures higher by 20 °C than those cultured in Matrigel, taking the 3D spheroid configuration (Figure 7A). Spheroids needed lower temperatures than monolayers to reach the same ablation percentage at all applied ultrasound intensities. Add to that, the correlation between the temperature and percentage of tumor ablation showed less scattering with spheroids in comparison to monolayers of cells, emphasizing that spheroids are less sensitive to changes in temperature.

**Figure 7.**
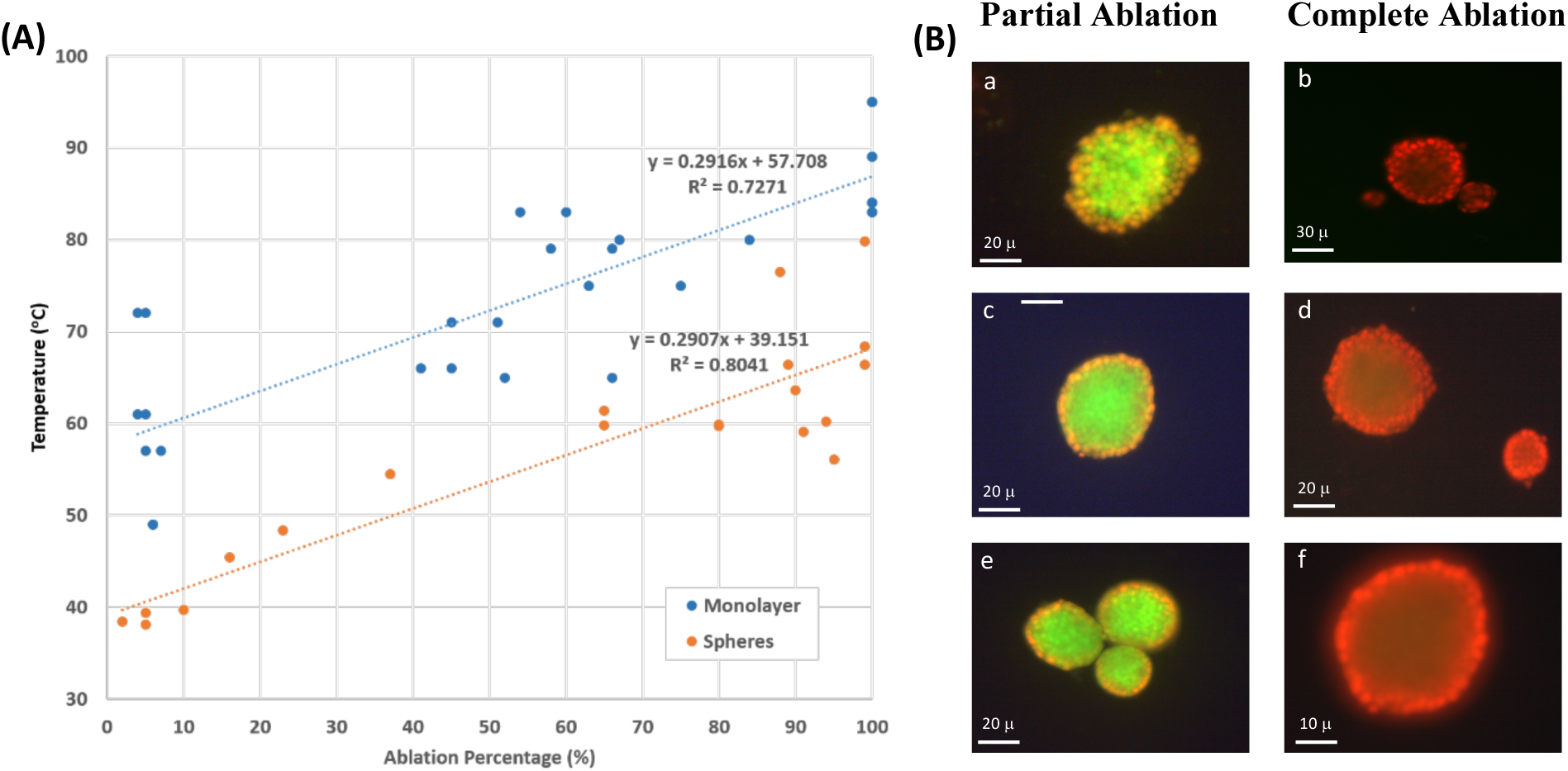
2D vs 3D culture ablation. (A) Final temperature versus ablation percentage of monolayers and spheroids for distinct combinations of DC and SD. (B) Microscope images of spheroids post sonication. With partial ablation, the outmost surface showed red stains (dead cells), whereas the cells closer to the center remained alive. With complete ablation, the red color dominates.

Images using Fluorescent Cyto3D Live/Dead Assay of spheroids post ultrasound sonication showed partial ablation of cultured cells, described by the death of the outermost layer (stained in red) while the innermost layer remained intact (stained in green), when sonicating at low values of DC. However, at the end of ultrasound sonication with higher DC and 𝐼*_SPPA_* complete ablation of spheroids was achieved (Figure 7B), where most cells in each cluster, whether in the outermost layer or the innermost one, became fully ablated after absorbing more power per unit area.

### 3.4 Numerical Simulations

In this section, the obtained experimental and numerical results are presented for the different operating conditions of DC and transducer pressure levels to analyze their effect on the ablation rate for the different cell culture types (monolayer and spheroids).

The developed mathematical models were first validated against the experimental data for both cell cultures. To that end, the model was simulated using the geometry of the adopted transducer in the experiment with a focal distance of 51.4 mm, an external diameter of 33 mm, and a fundamental frequency of 2 MHz. Our computational model describing the ablation of 2D and 3D cultured cells showed agreement with the experimental results in terms of the maximum temperature reached at the focal region within the cell cultures with a maximum error of 15 % as shown in Table 2, which is deemed acceptable in practice.

**Table 2.**
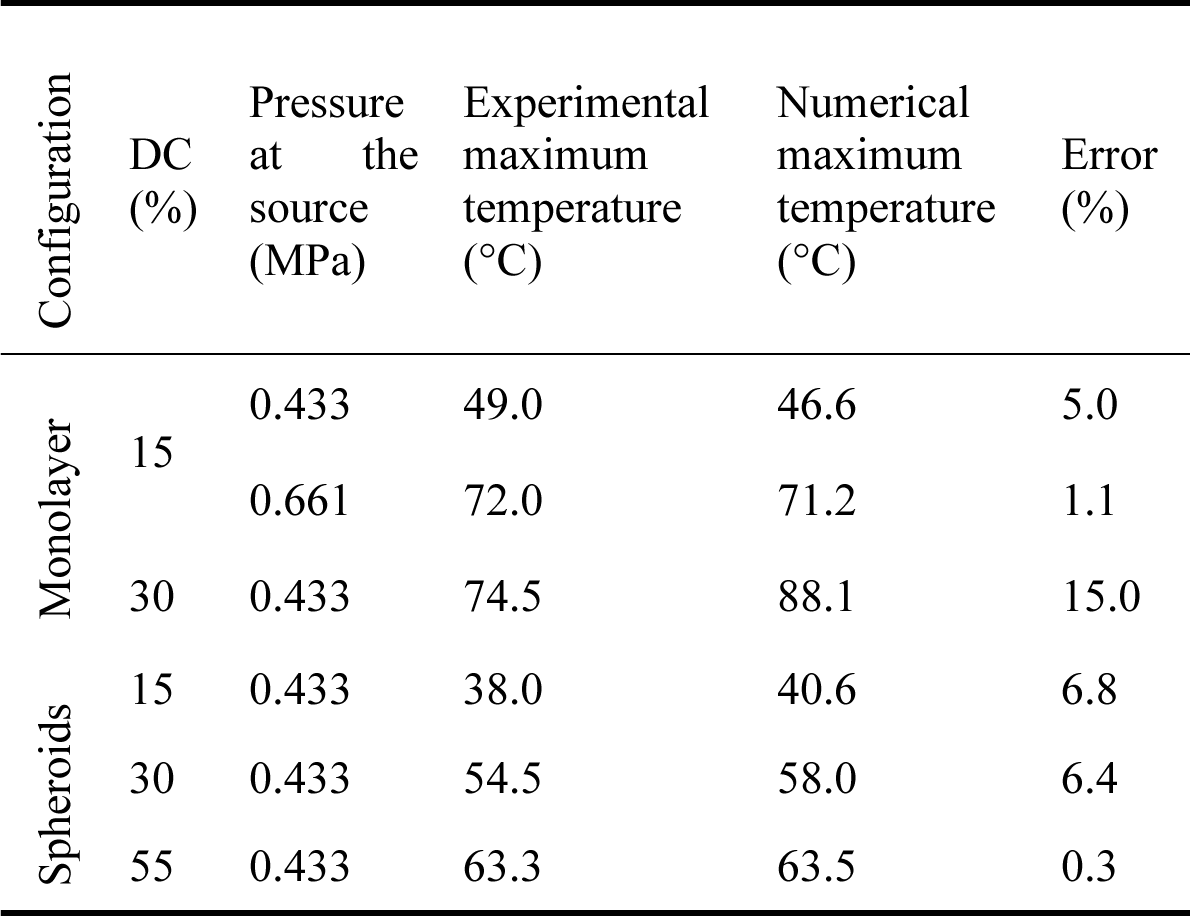
Comparison between the experimental and numerical results.

| Configuration | DC<br>(%) | Pressure<br>at the<br>source<br>(MPa) | Experimental<br>maximum<br>temperature<br>(°C) | Numerical<br>maximum<br>temperature<br>(°C) | Error<br>(%) |
| --- | --- | --- | --- | --- | --- |
| Monolayer | 15 | 0.433 | 49.0 | 46.6 | 5.0 |
|  |  | 0.661 | 72.0 | 71.2 | 1.1 |
|  | 30 | 0.433 | 74.5 | 88.1 | 15.0 |
| Spheroids | 15 | 0.433 | 38.0 | 40.6 | 6.8 |
|  | 30 | 0.433 | 54.5 | 58.0 | 6.4 |
|  | 55 | 0.433 | 63.3 | 63.5 | 0.3 |

In both the monolayer and spheroid simulations, surface plots spanning one quarter of the disk demonstrated a uniform distribution of thermal energy at low pressure runs of 0.433 MPa, with a minimal temperature gradient (less than 0.1 °C) between the focal region and the remainder of the culture medium. This was achieved with both low and high DC values (Figure 8A&B). However, at high pressure runs of 0.661 MPa, a large temperature gradient was recorded between the focal region and its immediate surrounding reaching up to 21.7 °C, computed at the 55 % high DC in the monolayer configuration, and up to 14 °C difference computed at a DC of 30 % with the spheroid configuration (Table 3). Nevertheless, within the focal region, the temperature distribution was relatively uniform with a negligible temperature gradient.

**Figure 8.**
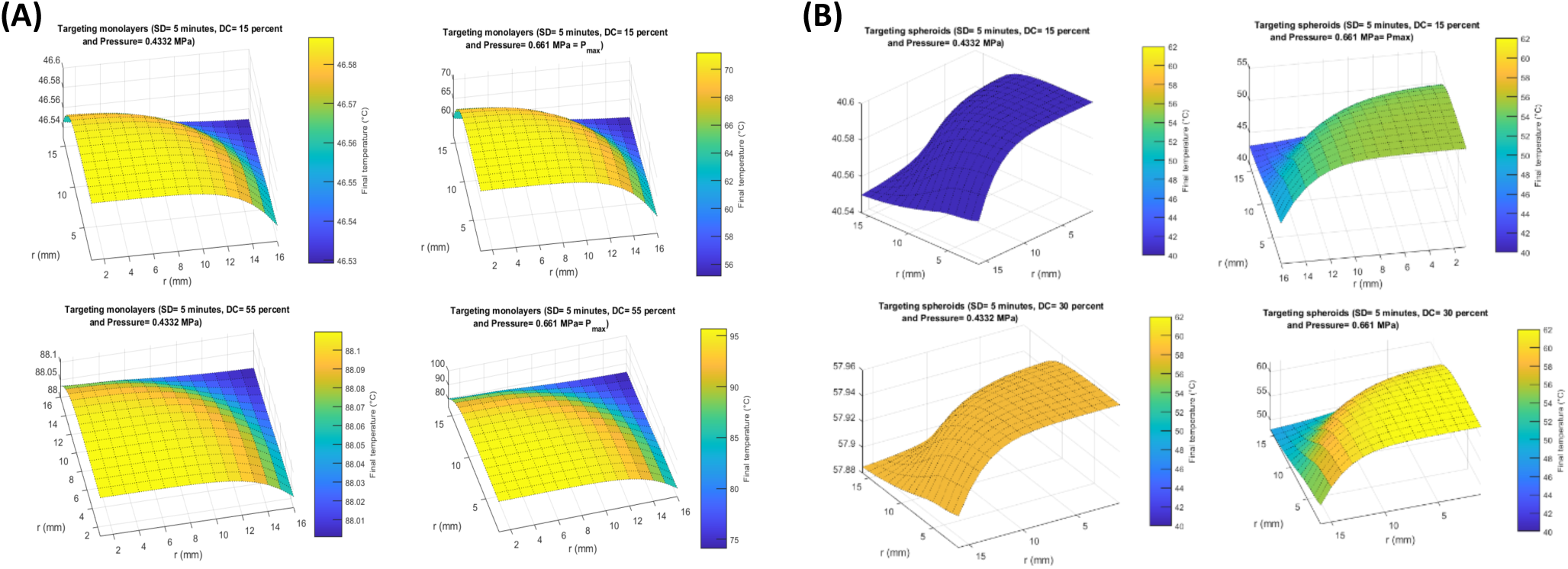
Range of final temperatures (°C) in the plane passing through the cells modeled as (A) monolayers and (B) spheroids at a DC of 15 % and 30 %, and a pressure of 0.433 MPa and 0.661 MPa after the simulation of ultrasound sonication for 5 minutes.

**Table 3.** Final temperature range.

| Configuration | DC (%) | Pressure at the source (MPa) | Final Temperature Range (°C) |
| --- | --- | --- | --- |
| Monolayer | 15 | 0.433 | [46.53; 46.59] |
|  |  | 0.661 | [55.13; 71.21] |
|  | 30 | 0.433 | [88.00; 88.11] |
|  |  | 0.661 | [74.01; 95.71] |
| Spheroids | 15 | 0.433 | [46.53; 46.59] |
|  |  | 0.661 | [55.13; 71.21] |
|  | 30 | 0.433 | [88.00; 88.11] |
|  |  | 0.661 | [74.01; 95.71] |

The configuration of the cultured cells did not affect how the latter interacted with the thermal energy induced by ultrasound sonication. Both monolayer and spheroid cell cultures showed similar behaviors regarding thermal responses to the sonication conditions. However, higher temperature values were attained by the monolayer culture compared to the spheroids for the same sonication parameters. As was demonstrated in the experimental work, a significant difference of up to 20 °C was recorded between the temperature reached by monolayers and that by spheroids. This difference in temperature was more apparent as DC went up.

## 4. Discussion

High intensity focused ultrasound has been widely used as a novel technology for cancer treatment due to its minimal invasiveness, cost-effectiveness, and high precision (44). Our research focused on the effect of pulsed HIFU on 2D and 3D breast cancer cell culture models to explore the effect of cell configuration and ultrasound parameters on the efficiency of HIFU for ablating breast cancer cells. A HIFU treatment was considered efficient if it led to an ablation percentage greater than or equal to 90 %. Our investigation delved comprehensively into the thermal ablation of 2D and 3D cancer cell cultures across a spectrum of ultrasound intensities, duty cycles, and sonication durations. Key metrics of interest included the percentage of cellular ablation and the maximum temperature reached during ablation. By assessing the response of these two culture models to ultrasound sonication, we aimed to characterize the optimal ultrasound parameters necessary for effective cancer cell ablation. Furthermore, we developed a mathematical model to elucidate how cells in monolayer and spheroid cultures respond to pulsed HIFU. These simulations were subsequently compared with experimental data to validate the model’s accuracy. This integrated approach allowed us to predict cellular ablation outcomes under varying conditions, encompassing alterations in duty cycles and maximum pressure levels, further enhancing our understanding of HIFU’s potential for cancer treatment.

### 4.1 Duty Cycle Affects Degree of Cellular Ablation

The duty cycle in ultrasound sonication denotes the fraction of the burst period during which ultrasound amplitude is nonzero, referred to as the ON region. A higher DC corresponds to an extended duration in which cells can absorb energy, resulting in a higher energy input per unit area. Conversely, a lower DC corresponds to a shorter ON region and a more extended OFF region, during which the absorbed energy dissipates into the surrounding control volume encompassing the target.

In both monolayer and spheroidal configurations, the percentage of ablated tumor cells was highly influenced by the DC of ultrasound waves, which increased as a function of increasing DC. When targeting bovine liver tissue in vitro with a 40 kHz frequency difference between the inner and outer loops of a dual-frequency transducer at 160 W ultrasonic power, Zhu et al. (45) reported a significant change in the structural form and dimensions of the lesion as DC was changed. With low DC values, ranging between 5-20 %, no coagulation necrosis was recorded, unlike with a DC of 30 % and above where coagulation necrosis appeared, with the highest ablation percentage occurring at a DC of 50 %. Similarly, our study showed that a low DC of 15 % was totally inefficient, while a DC of 55 % caused the ablation of more than 90 % of the tumor cells. Increasing DC elongated the heating time, and diminished the dissipation time, characterized by a shorter inactive treatment interval, allowing thermal ablation to dominate as more shock wave arrays continued to hit the targeted tissue (46). Indeed, higher temperatures were attained with higher DC values as a result of longer heat depositing intervals. However, there is a potential trade-off. Increased ultrasonic pulse frequency, driven by higher DC values, elevates the risk of inducing mechanical disintegration and localized fragmentation of the targeted tissue due to inertial cavitation, as noted in prior research (10). After a certain threshold, even with further increases in DC values beyond those employed in our study, a transition occurs wherein the efficacy of thermal ablation begins to diminish, yielding to the predominance of mechanical ablation as the more efficient process.

Our findings revealed that applying ultrasound sonication for durations of 5 minutes or 10 minutes yielded nearly similar ablation percentages, with negligible differences, particularly at low and high DC values for both monolayers and spheroids. This observation led to the hypothesis that cells might reach a state of equilibrium in terms of power absorption and energy dissipation before the conclusion of the sonication session, indicating a saturation in behavior at an optimal exposure duration. Consequently, a 5-minute treatment duration may be considered more efficient, as it achieves the desired ablation percentages with less time and energy expenditure.

### 4.2 Monolayer vs. Spheroid Ablation Temperature

Cells modeled as spheroids were ablated at a threshold temperature significantly lower than that of monolayers (approximately 20 °C less), demonstrating less sensitivity of spheroids than monolayers to thermal ablation. Both cell configurations differed in oxygen and nutrient availability and distribution among the cells in each cluster. During treatment, the cells absorbed the acoustic intensity in the ON region for further energy dissipation to the surroundings during the OFF region. In general, spheroids have a cell growth medium with a higher thermal conductivity than that of monolayers. They also have a higher surface-to-volume ratio. Thus, cells in the innermost core of the spheroids would dissipate power into the surrounding cells during the OFF region, unlike cells in monolayers where the dissipated power would mostly spread across to the surrounding cell growth medium.

Results showed a uniform distribution of heat across the focal region in both cell configurations. However, approximately 12.5 % higher maximum intensity was recorded at the same applied pressure in spheroids than in monolayers, which increased as the applied pressure was also increased (47). This can potentially be due to an additional absorption coefficient and specific heat capacity offered by the hydrogel to spheroids (Table 1), causing the latter to absorb much more heat than the hydrogel-lacking monolayers, while only leading to slow increase in temperature. This could explain why higher intensities causing higher temperatures were required to implement the same ablation percentage in the monolayer case. For the same sonication parameters, however, the spheroids significantly heated up to lower temperatures, implying they were more susceptible to mechanical ablation than thermal ablation at this stage compared to monolayers due to microenvironment disruption altering tumor cell viability (48). With high negative pressures and shorter HIFU pulses (49), residual gas bubbles present within the spheroids produced cavitating bubbles causing mechanical damage rather than thermal damage in a series of cavitation events taking place (50, 51).

### 4.3 Partial and Complete Cellular Ablation in Spheroids

Cells at the outermost layer of the spheroids got ablated before those in the innermost layer which remained intact at low DC and sonication intensities. As DC and intensity were increased, full ablation of all the cells in the clusters was achieved at the end of each sonication session.

Spheroids have an oxygen concentration gradient from the outer surface to the inner core, with cells located on the outermost surface of the cluster being exposed to oxygen more than those at the core which became hypoxic spots (20). Hence, the latter were more susceptible to ablation at higher intensities while reaching higher temperature levels. Nonetheless, microscopic images showed that the outer surface of a cell cluster got ablated first, followed by the hypoxic core, suggesting a gradient in the distribution of ultrasound power per unit volume while being transferred across the cells. In general, 3D cell cultures have a more intricate interconnection between cells, resulting in the formation of gradients in pH, oxygen concentration, and metabolic activity (52). These cell-cell interactions could form a shielding effect to the cells present at the core despite the uniform thermal distribution across.

### 4.4 Model Validation

Improving the effectiveness and efficiency of HIFU treatment requires simulating the way non-linear acoustic waves travel through various layers of media using an ultrasound transducer. These simulations involve trying out different strategies for focusing the waves and observing how they result in an increase in the temperature of the targeted tissue, leading to the formation of lesions. These computational studies are vital for planning and optimizing HIFU procedures directed for clinical use.

Our computational model highlighted the difference in the maximal temperature reached by both simulated cell configurations. In specific, spheroids were subjected to a higher heat deposition due to their higher acoustic impedance (higher density of the spheroids culture media (RPMI) compared to that of the monolayer (DMEM)). Moreover, spheroids in general are characterized by a thicker layer (117 μm vs 15 μm), which increases their total impedance. For these reasons, higher acoustic energy was deposited in the cells, resulting in higher heat fixation. Nonetheless, the temperature of the spheroids was significantly lower even though both culture media had similar specific heat capacities. This could be attributed to the 75 % higher conductivity of spheroids gel (0.53 W/K.m) compared to that of the monolayer (0.13 W/K.m). This resulted in higher heat conduction and dissipation to the surroundings, leading to reduced temperature levels recorded with the spheroids. This effect, combined with the additional specific heat capacity from the hydrogel medium, contributes to a slower increase in temperature within the spheroids compared to monolayers.

Furthermore, at low DC, the spheroids were ablated at the boundary first while the inner cells remained intact. Upon the increase in DC, all the cells were ablated including the innermost ones. Our model showed that the temperature of the spheroids was uniform throughout the culture medium with no significant temperature gradient within the focal region. Similar observations were obtained by Zhou et al. (53), where the center of the ablated cancer appeared analogous to viable cells after H&E staining. These cells maintained their characteristics of cytologic staining and nuclear chromatin without any signs of breakdown. However, electron microscopy revealed that the cytoplasm of those normal-appearing cancer cells contained vacuoles, where the cell membranes were disintegrated with unidentified organelle structures, suggesting an irreversible cell death with the preservation of cellular structure induced by thermal fixation instead of incomplete coagulation necrosis (53). Add to that, the central part of the ablated tumor resisted degradation since the wound-healing process could not extend to this region right after HIFU treatment. In contrast, and in the peripheral region, cancer cells had the typical characteristics of lethal and irreversible cell damage as coagulation necrosis. Therefore, NADH-diaphorase stain was found more accurate and objective than H&E staining in assessing acute cell death because it is based on the presence or absence of enzyme function instead of changes in the cellular structure. These conclusions were also emphasized by Wang et al. (54) where the peripheral region was lethally damaged while the central region, which was thermally fixed, looked normal and similar to viable cells with the preservation of cell structure. Hence, we should check if the adopted staining techniques are indeed suitable to accurately reflect the status of cell viability.

However, to explain such a phenomenon, assuming the staining observations were reliable, one possible reason that could be attributed to such findings is the effect of mechanical ablation instead of thermal ablation at this level with the application of pulsed HIFU. Mechanical effects could result in the ablation of spheroids at lower temperatures compared to monolayers. In fact, using high intensity ultrasound waves, mechanical effects such as cavitation and microstreaming are dominant (55). Pulsed HIFU can induce bubble formation that disrupts the tissues and causes its liquefaction after the bubble is exploded by subsequent acoustic waves. Such mechanical effects are more prominent when pulsed HIFU beams are used instead of continuous HIFU (56), where the burst duration or pulse length is shorter than the time needed for the cells to boil, form a bubble, and then burst (57). Accordingly, bubbles that form on the inner or central region could be protected by the cell-cell and cell-matrix interactions, thus resisting HIFU ablation more than the outermost cells where the bubbles could be easily depleted.

The formation of microbubbles in spheroids could also explain the difference in temperature levels attained in this 3D cell culture compared to the monolayer. Also, the bubbles formed by HIFU could potentially create a shielding layer that prevents the HIFU wave penetration into the core, causing the waves’ reflection and backscatter, thereby reducing the amount of thermal energy dissipated in the cells (58, 59). Such acoustic wave reflection could not be captured by the adopted acoustic wave propagation model since it neglects the formation of the bubbles. Moreover, the formation of these bubbles may be limited in the monolayer culture due to its 2D geometry and small thickness compared to the more realistic 3D spheroid culture.

## 5. Conclusion

High intensity focused ultrasound is gaining fast clinical acceptance for thermal ablation of cancerous tumors. Studying the effect of HIFU parameters on 2D and 3D models of tumor cells is essential for determining the best set of parameters to depict the most significant outcomes of ablation area, temperature rise and tumor damage without any disease recurrence. Spheroid cell culturing resembled *in vivo* tumors more than the monolayer cell culture formation and reached lower temperature elevation at similar acoustic intensities and duty cycles. Unlike monolayers, spheroids could allow for the interplay of both thermal and mechanical ablation due to their extracellular-cell-medium mimicking core as DC increased. The ultrasound DC affected the degree of tumor ablation whereby an increase in DC resulted in an increase in the ablation percentage. However, ultrasonic sonication time had little effect on the ablation percentage. Numerical simulations of both culture configurations emphasized a uniform distribution of heat across the cultured cells. At high DC and spatial-peak pulse-average intensity, complete ablation of spheroids took place, yet at the lower end only the outmost layer got ablated. Pulsed HIFU could cause mechanical ablation of cells through cavitation dominating thermal ablation. Our study had some limitations whereby all interactions of cells with the extracellular medium as in an *in vivo* setting were not taken into consideration. Temperature measurements were highly dependent on the sensitivity of the thermocouple used and its right positioning in the focal region without damaging the cultured medium. Add to that, the same petri dish contained varying sizes of spheres, posing challenges in examining how HIFU parameters affected spheroid dimensions. We were also unable to quantify the number of cells before subjecting them to sonication. Nonetheless, our work emphasized the ability of HIFU to ablate 2D and 3D cultured tumors while identifying the effect of ultrasound parameters on the ablation percentage, area of damage and temperature rise post sonication.

